# CAT – A Computational Anatomy Toolbox for the Analysis of Structural MRI Data

**DOI:** 10.1101/2022.06.11.495736

**Authors:** Christian Gaser, Robert Dahnke, Paul M Thompson, Florian Kurth, Eileen Luders, Alzheimer’s Disease Neuroimaging Initiative

**Affiliations:** Department of Psychiatry and Psychotherapy, Jena University Hospital, Jena, Germany; Department of Neurology, Jena University Hospital, Jena, Germany; German Center for Mental Health (DZPG); Imaging Genetics Center, Stevens Neuroimaging & Informatics Institute, Keck School of Medicine, University of Southern California, Los Angeles, CA, USA; School of Psychology, University of Auckland, Auckland, New Zealand; Department of Women’s and Children’s Health, Uppsala University, Uppsala, Sweden

**Keywords:** brain, computational anatomy, longitudinal, morphometry, SPM12, CAT12, MRI, ROI, VBM, cortical thickness, cortical surface, cortical folding, Alzheimer’s disease

## Abstract

A large range of sophisticated brain image analysis tools have been developed by the neuroscience community, greatly advancing the field of human brain mapping. Here we introduce the *Computational Anatomy Toolbox* (CAT) – a powerful suite of tools for brain morphometric analyses with an intuitive graphical user interface, but also usable as a shell script. CAT is suitable for beginners, casual users, experts, and developers alike providing a comprehensive set of analysis options, workflows, and integrated pipelines. The available analysis streams – illustrated on an example dataset – allow for voxel-based, surface-based, as well as region-based morphometric analyses. Notably, CAT incorporates multiple quality control options and covers the entire analysis workflow, including the preprocessing of cross-sectional and longitudinal data, statistical analysis, and the visualization of results. The overarching aim of this article is to provide a complete description and evaluation of CAT, while offering a citable standard for the neuroscience community.

## Main

The brain is the most complex organ of the human body, and no two brains are alike. The study of the human brain is still in its infancy, but rapid technical advances in image acquisition and processing are enabling ever more refined characterizations of its micro- and macro-structure. Enormous efforts, for example, have been made to map differences between groups (e.g., young vs. old, diseased vs. healthy, male vs. female), to capture changes over time (e.g., from infancy to old age, in the framework of neuroplasticity, as a result of a clinical intervention), or to assess correlations of brain attributes (e.g., measures of length, volume, shape) with behavioral, cognitive, or clinical parameters. Popular neuroimaging software packages include tools for analysis and visualization, such as SPM (https://www.fil.ion.ucl.ac.uk/spm), FreeSurfer (https://surfer.nmr.mgh.harvard.edu), the Human Connectome Workbench (https://www.humanconnectome.org/software/connectome-workbench), FSL (https://www.fmrib.ox.ac.uk/fsl), BrainVISA (http://www.brainvisa.info), CIVET (https://mcin.ca/technology/civet), or the LONI tools (https://www.loni.usc.edu/research/software), just to name a few.

SPM (short for *Statistical Parametric Mapping*) is one of the most frequently used software packages. Its library of accessible and editable scripts provide an ideal basis to extend the repertoire of preprocessing and analysis options. Over the years, SPM has inspired developers to create powerful tools that use SPM’s functionality and interface (https://www.fil.ion.ucl.ac.uk/spm/ext), but that are more than mere extensions of SPM offering a comprehensive range of cutting-edge preprocessing and analysis options across the whole analysis spectrum, from the initial data processing to the final visualization of the statistical effects.

One such tool is CAT (short for *Computational Anatomy Toolbox;* https://neuro-jena.github.io/cat). CAT constitutes a significant step forward in the field of human brain mapping by adding sophisticated methods to process and analyze structural brain MRI data using voxel-, surface-, and region-based approaches. CAT is available as a collection of accessible scripts, with an intuitive user interface, and uses the same batch editor as SPM, which allows for a seamless integration with SPM workflows and other toolboxes, such as Brainstorm (Tadel et al., 2011) and ExploreASL (Mutsaerts et al., 2020). Not only does this enable beginners and experts to run complex state-of-the-art structural image analyses within the SPM environment, it will also provide advanced users as well as developers the much appreciated option to incorporate a wide range of functions in their own customized workflows and pipelines.

## Results

### Concept of CAT

CAT12 is the current version of the CAT software and runs in Matlab (Mathworks, Natick, MA) or as a standalone version with no need for a Matlab license. It was originally designed to work with SPM12 (http://www.fil.ion.ucl.ac.uk/spm/software/spm12) and is compatible with Matlab versions 7.4 (R2007a) and later. No additional software or toolbox is required. The latest version of CAT can be downloaded here: https://neuro-jena.github.io/cat. The pre-compiled standalone version for Windows, Mac, or Linux operating systems can be downloaded here: https://neuro-jena.github.io/enigma-cat12/#standalone. All steps necessary to install and run CAT are documented in the user manual (https://neuro-jena.github.io/cat12-help) and in the complementary online help, which can be accessed directly via CAT’s help functions. The CAT software is free but copyrighted and distributed under the terms of the GNU General Public License, as published by the Free Software Foundation.

CAT can be either started through SPM, from the Matlab command window, from a shell, or as a standalone version. Except when called from the command shell (CAT is fully scriptable), a user interface will appear (see **Figure 1**) allowing easy access to all analysis options and most additional functions. In addition, a graphical output window will display the interactive help to get started. This interactive help will be replaced by the results of the analyses (i.e., in that same window), but can always be called again via the user interface.

**Figure 1:**
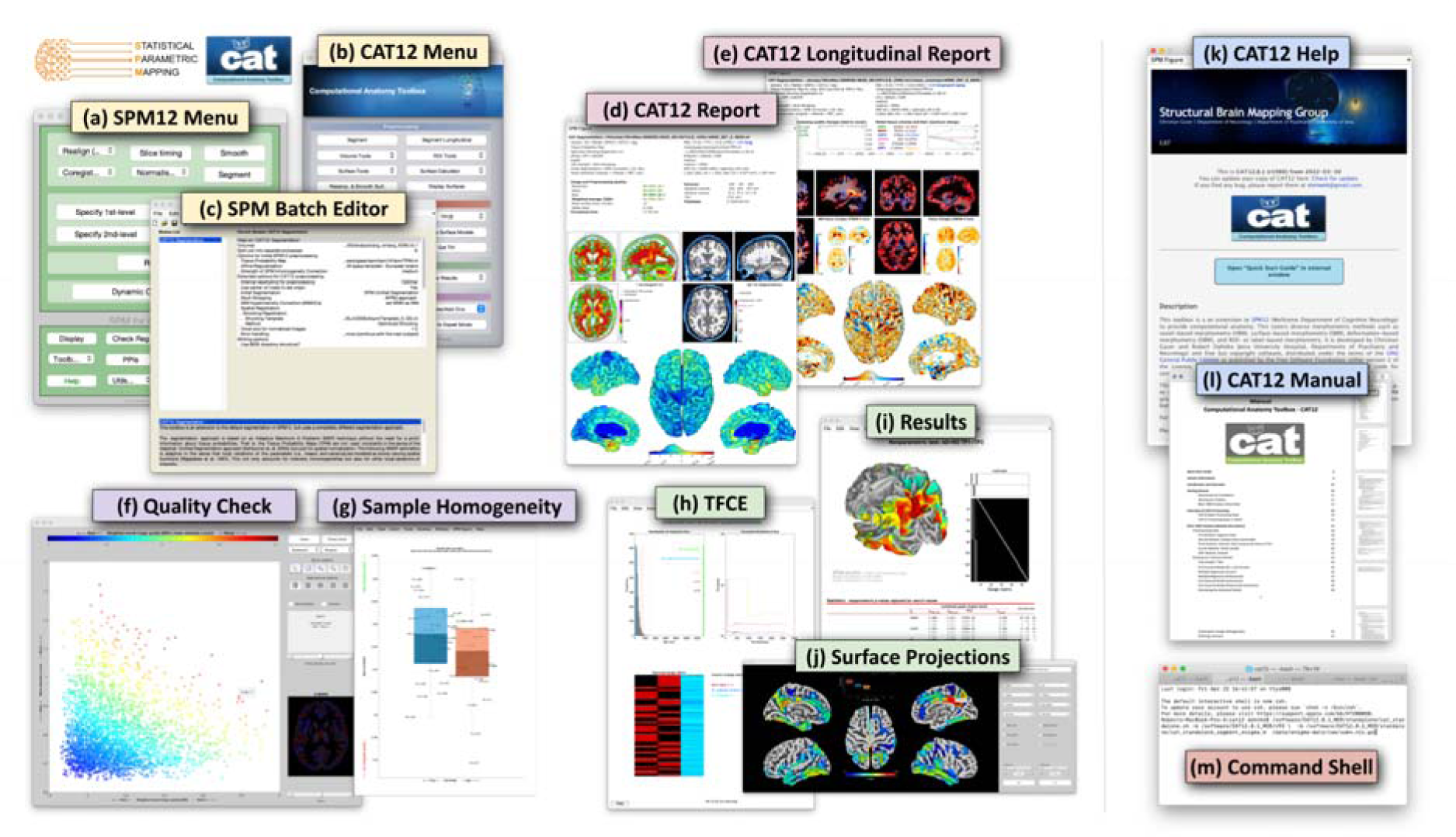
Elements of the graphical user interface. The SPM menu (a) and CAT menu (b) allow access to the (c) SPM batch editor to control and combine a variety of functions. At the end of the processing stream, cross-sectional and longitudinal outputs are summarized in a brain-specific one-page report (d, e). In addition, CAT provides options to check image quality (f) and sample homogeneity (g) to allow outliers to be removed before applying the final statistical analysis, including *threshold-free cluster enhancement* – TFCE (h); the numerical and graphical output can then be retrieved (i), including surface projections (j). For beginners, there is an interactive help (k) as well as a user manual (l). For experts, command line tools (m) are available under Linux and MacOS.

### Computational Morphometry

CAT’s processing pipeline (see **Figure 2**) contains two main streams: (1) voxel-based processing for voxel-based morphometry (VBM) and (2) surface-based processing for surface-based morphometry (SBM). The former is a prerequisite for the latter, but not the other way round. Both processing streams can be extended to include additional steps for (3) region-based processing and region-based morphometry (RBM).

**Figure 2:**
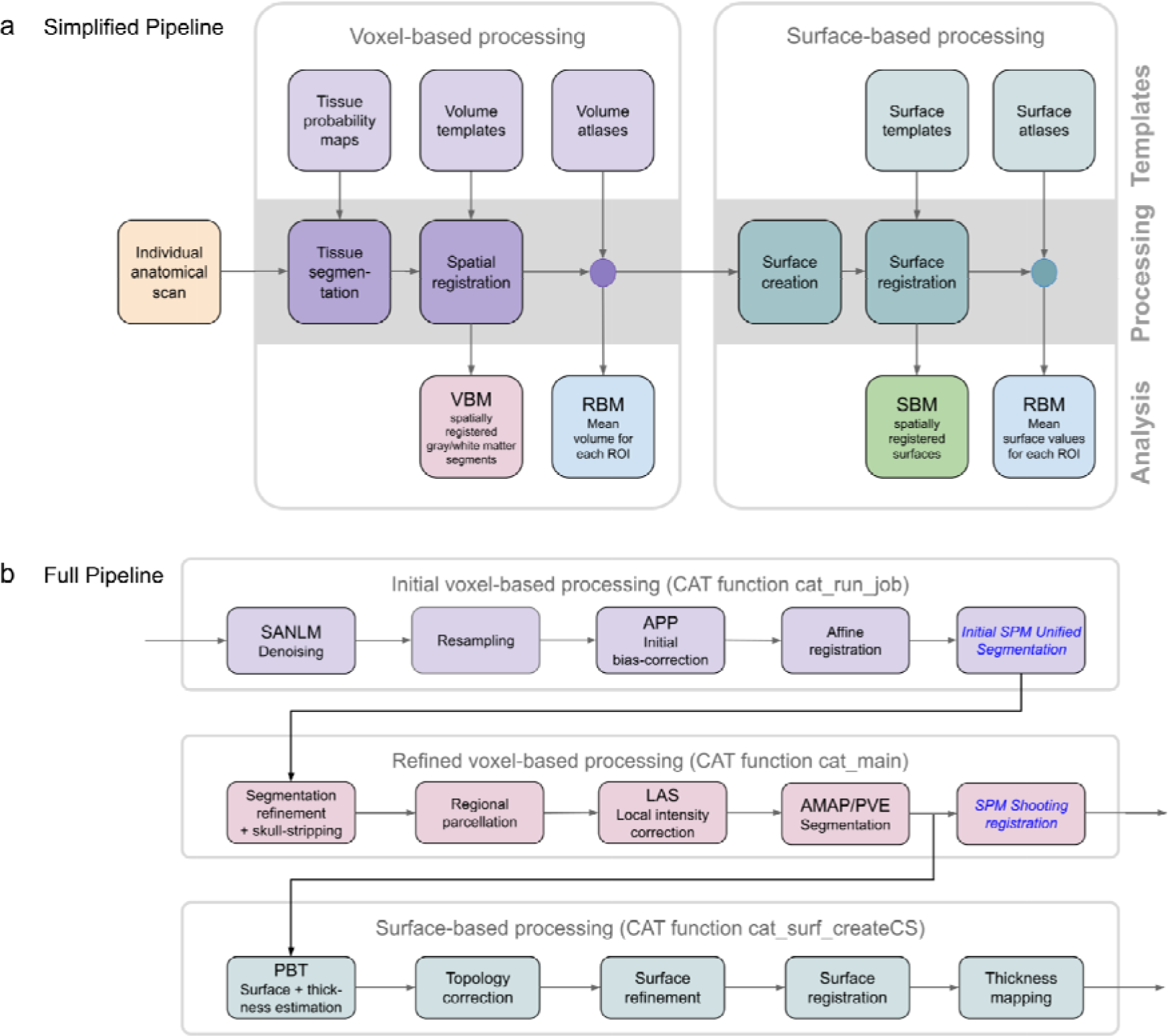
Main processing streams. (a) Simplified pipeline: Image processing in CAT can be separated into a mandatory voxel-based processing stream and an optional subsequent surface-based processing stream. Each stream requires different templates and atlases and, in addition, tissue probability maps for the voxel-based stream. The voxel-based stream consists of two main modules – for tissue segmentation and spatial registration – resulting in spatially registered (and modulated) gray matter / white matter segments, which provides the basis for *voxel-based morphometry* (VBM). The surface-based stream also consists of two main modules – for surface creation and registration – resulting in spatially registered surface maps, which provide the basis for *surface-based morphometry* (SBM). Both streams also include an optional module each to analyze *regions of interest* (ROIs) resulting in ROI-specific mean volumes (mean surface values, respectively). This provides the basis for *region-based morphometry* (RBM). (b) Detailed pipeline: To illustrate the differences from SPM, the CAT pipeline is detailed with its individual processing steps. The SPM methods used are shown in blue and italic font: images are first denoised by spatially adaptive non-local means (SANLM) filter (Manjón et al., 2010) and resampled to an isotropic voxel size. After applying an initial bias correction to facilitate the affine registration, SPM’s unified segmentatio (Ashburner & Friston, 2005) is used for the skull stripping and as a starting estimate for the adaptive maximum a posteriori (AMAP) segmentation (Rajapakse et al., 1997) with partial volume estimation (PVE) (Tohka et al., 2004). In addition, SPM’s segmentation is used to locally correct image intensities. Finally, the outcomes of the AMAP segmentation are registered to the MNI template using SPM’s shooting registration. The outcomes of the AMAP segmentation are also used to estimate cortical thickness and the central surface using a projection-based thickness (PBT) method (Dahnke et al., 2013). More specifically, after repairing topology defects (Yotter, Dahnke, et al., 2011) central, pial and white matter surface meshes are generated. The individual left and right central surfaces are then registered to the corresponding hemisphere of the FreeSurfer template using a 2D version of the DARTEL approach (Ashburner, 2007). In the final step, the pial and white matter surfaces are used to refine the initial cortical thickness estimate using the FreeSurfer thickness metric (Fischl & Dale, 2000; Masouleh *et al., 2020)*.

### Voxel-based Processing

Voxel-based processing steps can be roughly divided into a module for tissue segmentation, followed by a module for spatial registration.

⍰ Tissue Segmentation: The process is initiated by applying a *spatially adaptive non-local means* (SANLM) denoising filter (Manjón et al., 2010), followed by SPM’s standard *unified segmentation* (Ashburner & Friston, 2005). The resulting output serves as a starting point for further optimizations and CAT’s tissue segmentation steps: first, the brain is parcellated into the left and right hemispheres, subcortical areas, ventricles, and cerebellum. In addition, local white matter hyperintensities are detected (to be later accounted for during the spatial registration and the optional surface processing). Second, a local intensity transformation is performed to reduce effects of higher gray matter intensities in the motor cortex, basal ganglia, and occipital lobe. Third, an *adaptive maximum a posteriori* (AMAP) segmentation is applied which does not require any *a priori* information on the tissue probabilities (Rajapakse et al., 1997). The AMAP segmentation also includes a *partial volume estimation* (Tohka et al., 2004). **Figure 3** provides information on the accuracy of CAT’s tissue segmentation.
⍰ Spatial Registration: Geodesic Shooting (Ashburner & Friston, 2011) is used to register the individual tissue segments to standardized templates in the ICBM 2009c Nonlinear Asymmetric space (*MNI152NLin2009cAsym*; https://www.bic.mni.mcgill.ca/ServicesAtlases/ICBM152NLin2009), hereafter referred to as MNI space. While MNI space is also used in many other software packages, enabling cross-study comparisons, users may also choose to use their own templates. **Figure 3** provides information on the accuracy of CAT’s spatial registration.

**Figure 3:**
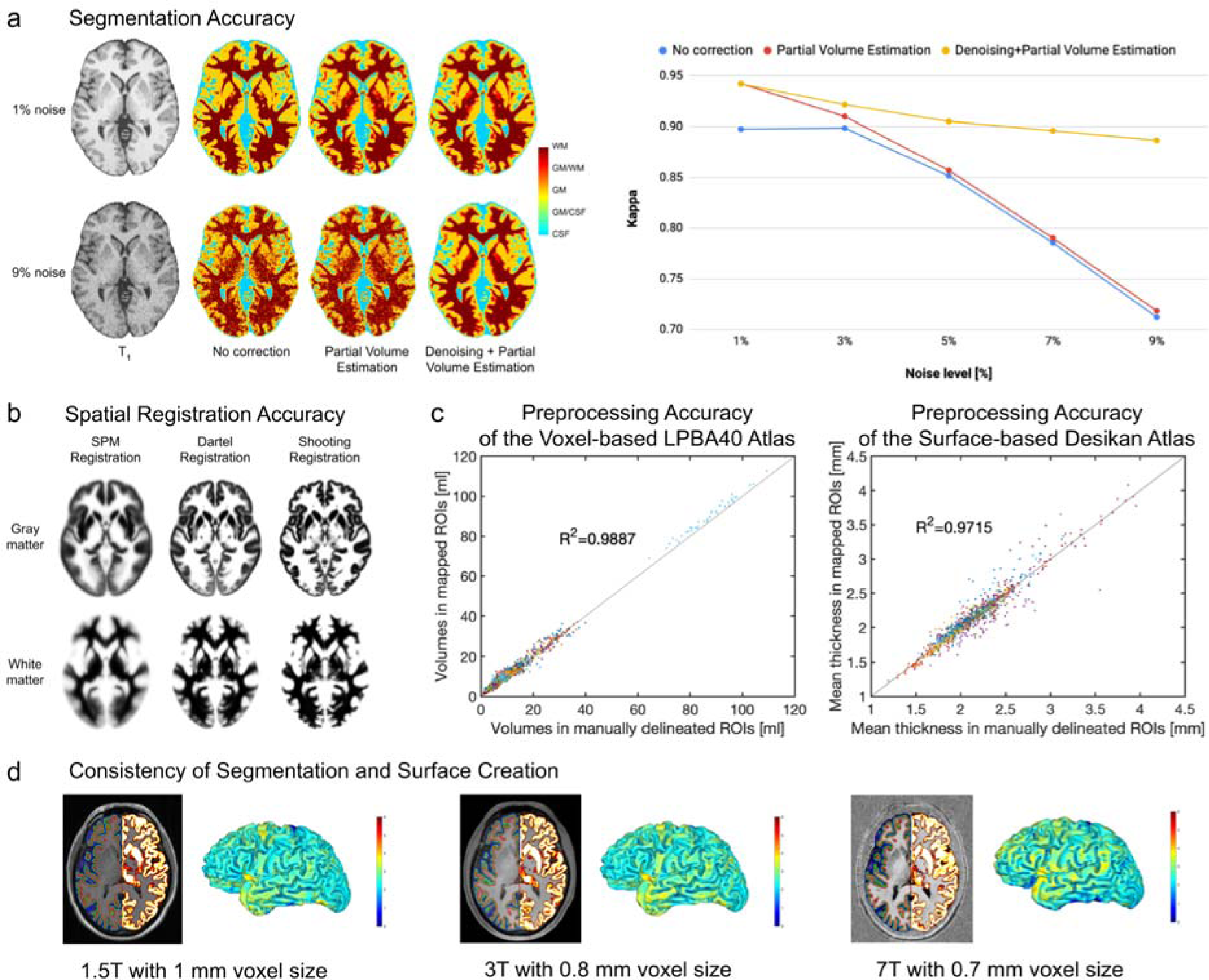
Evaluation of segmentation and registration accuracy. (a) *Segmentation Accuracy:* Most approaches for brain segmentation assume that each voxel belongs to particular tissue class, such as *gray matter* (GM), *white matter* (WM), or *cerebrospinal fluid* (CSF). However, the spatial resolution of brain images is limited, leading to so-called *partial volume effects* (PVE) in voxels containing a mixture of different tissue types, such as GM/WM and GM/CSF. As PVE approaches are highly susceptible to noise, we combined the PVE model (Tohka et al., 2004) with a spatial adaptive non-local means denoising filter (Manjón et al., 2010). To validate our method, we used a ground truth image from the BrainWeb (Aubert-Broche et al., 2006) database with varying noise levels of 1-9%. The segmentation accuracy for all tissue types (GM, WM, CSF) was determined by calculating a *kappa* coefficient (a kappa coefficient of 1 means that there is perfect correspondence between the segmentation result and the ground truth). *Left panel:* The effect of the PVE model and the denoising filter on the tissue segmentation at the extremes of 1% and 9% noise. *Right panel:* The kappa coefficient over the range of different noise levels. Both panels demonstrate the advantage of combining the PVE model with a spatial adaptive non-local means denoising filter, with particularly strong benefits for noisy data. (b) *Registration Accuracy:* To ensure an appropriate overlap of corresponding anatomical regions across brains, high-dimensional nonlinear spatial registration is required. CAT uses a sophisticated Shooting approac (Ashburner & Friston, 2011), together with an average template created from the IXI dataset (http://www.brain-development.org). The figure shows the improved accuracy (i.e., a more detailed average image) when spatially registering 555 brains using the so-called ‘shooting’ registration and the Dartel registration compared to the SPM standard registration. (c) *Preprocessing Accuracy:* We validated the performance of region-based morphometry (RBM) in CAT by comparing measures derived from automatically extracted regions of interest (ROI) versus manually labeled ROIs. For the voxel-based analysis, we used 56 structures, manually labeled in 40 brains that provided the basis for the LPBA40 atlas (Shattuck et al., 2007). The gray matter volumes from those manually labeled regions served as the ground truth against which the gray matter volumes calculated using CAT and the LPBA40 atlas were then compared. For the surface-based analysis, we used 34 structures that were manually labeled in 39 brains according to Desikan (Desikan et al., 2006). The mean cortical thickness from those manually labeled regions served as the ground truth against which the mean cortical thickness calculated using CAT and the Desikan atlas were compared. The diagrams show excellent overlap between manually and automatically labeled regions in both voxel-based (left) and surface-based (right) analyses. (d) *Consistency of Segmentation and Surface Creation:* Data from the same brain were acquired on MRI scanners with different isotropic spatial resolutions and different field strengths: 1.5T MPRAGE with 1 mm voxel size; 3T MPRAGE with 0.8 mm voxel size; and 7T MP2RAGE with 0.7 mm voxel size. Section views: The left hemispheres depict the central (green), pial (blue), and white matter (red) surfaces; the right hemispheres show the gray matter segments. Rendered Views: The color bar encodes point-wise cortical thickness projected onto the left hemisphere central surface. Both section views and hemisphere renderings demonstrate the consistency of the outcomes of the segmentation and surface creation procedures across different spatial resolutions and field strengths.

### Voxel-based Morphometry (VBM)

VBM is applied to investigate the volume (or local amount) of a specific tissue compartment (Ashburner & Friston, 2005; Kurth et al., 2015) - usually gray matter. VBM incorporates different processing steps: (a) tissue segmentation and (b) spatial registration as detailed above, and in addition (c) adjustments for volume changes due to the registration (modulation) as well as (d) convolution with a 3D Gaussian kernel (spatial smoothing). As a side note, the modulation step results in voxel-wise gray matter volumes that are the same as in native space (i.e., before spatial registration) and not corrected for brain size yet. To remove effects of brain size, users have at least two options: (1) calculating the *total intracranial volume* (TIV) and including TIV as a covariate in the statistical model (Malone et al., 2014) or (2) selecting ‘global scaling’ (see second level options in SPM). The latter is recommended if TIV is linked with (i.e., not orthogonal to) the effect of interest (e.g., sex), which can be tested (see ‘Design orthogonality’ in SPM).

### Surface-based Processing

The optional surface-based processing comprises a series of steps that can be roughly divided into a module for surface creation, followed by a module for surface registration.

⍰ Surface Creation: **Figure 3** illustrates the surface creation step in CAT for data obtained on scanners with different field strengths (1.5, 3.0, and 7.0 Tesla). CAT uses a projection-based thickness method (Dahnke et al., 2013) which estimates the initial cortical thickness and initial central surface in a combined step, while handling partial volume information, sulcal blurring, and sulcal asymmetries, without explicit sulcus reconstruction. After this initial step, topological defects (i.e., anatomically incorrect connections between gyri or sulci) are repaired using spherical harmonics (Yotter, Dahnke, et al., 2011). The topological correction is followed by a surface refinement, which results in the final central, pial and white surface meshes. In the last step, the final pial and white matter surfaces are used to refine the initial cortical thickness estimate using the FreeSurfer thickness metric (Fischl & Dale, 2000; Masouleh et al., 2020). Alternatively, the final central surface can be used to calculate metrics of cortical folding, as described under **Surface-based Morphometry**.
⍰ Surface Registration: The resulting individual central surfaces are registered to the corresponding hemisphere of the FreeSurfer *FsAverage* template (https://surfer.nmr.mgh.harvard.edu/fswiki/FsAverage). During this process, the individual central surfaces are spherically inflated with minimal distortions (Yotter, Thompson, et al., 2011) and a one-to-one mapping between the folding patterns of the individual and template spheres is created by a 2D-version of the DARTEL approach (Ashburner, 2007; Yotter, Ziegler, et al., 2011). **Figure 3** provides information on the accuracy of CAT’s surface registration.

### Surface-based Morphometry (SBM)

SBM can be used to investigate cortical thickness or various parameters of cortical folding. The measurement of ‘cortical thickness’ captures the width of the gray matter ribbon as the distance between its inner and outer boundary at thousands of points (see **Figure 4**). To obtain measurements of ‘cortical folding’ the user has a variety of options in CAT, ranging from *Gyrification* (Luders et al., 2006) to *Sulcal Depth* (van Essen, 2005) to *Cortical Complexity* (Yotter, Nenadic, et al., 2011) to the *Surface Ratio* (Toro et al., 2008), as explained and illustrated in **Figure 4**. Similar to VBM, SBM incorporates a series of different steps: (a) surface creation and (b) surface registration as detailed above, and (c) spatial smoothing. As a side note, since the measurements in native space are mapped directly to the template during the spatial registration, no additional modulation (as in VBM) is needed to preserve the individual differences. In contrast to VBM, SBM does not require brain size corrections because cortical thickness and cortical folding are not closely associated with total brain volume (unlike gray matter volume) (Barnes et al., 2010).

**Figure 4:**
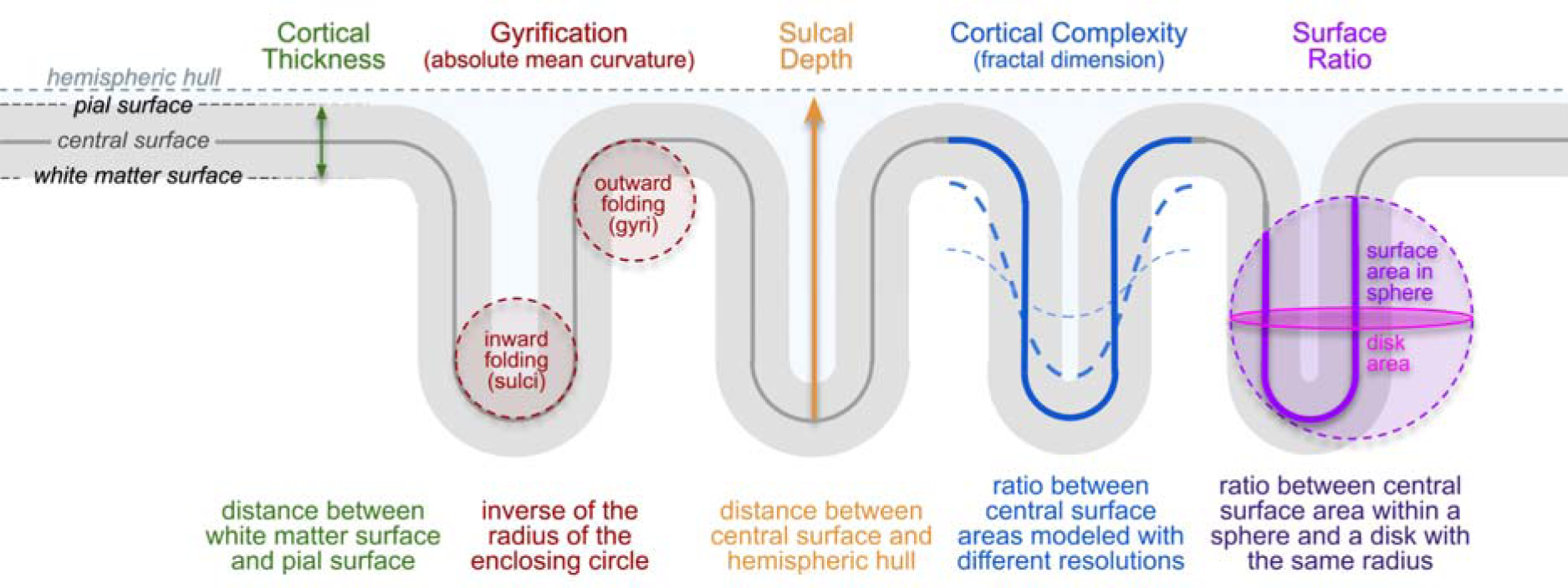
Cortical Measurements. Surface-based morphometry is applied to investigate cortical surface features (i.e., cortical thickness and various parameters of cortical folding) at thousands of surface points. *Cortical Thickness*: One of the best known and most frequently used morphometric measures is cortical thickness, which captures the width of the gray matter ribbon as the distance between its inner boundary (white matter surface) and outer boundary (pial surface). *Cortical Folding*: CAT provides distinct cortical folding measures, derived from the geometry of the central surface: ‘Gyrification’ is calculated via the absolute mean curvature (Luders et al., 2006) of the central surface. ‘Sulcal Depth’ is calculated as the distance from the central surface to the enclosing hull (van Essen, 2005). ‘Cortical Complexity’ is calculated using the fractal dimension of the central surface area from spherical harmonic reconstructions (Yotter, Nenadic, et al., 2011). Finally, ‘Surface Ratio’ is calculated as the ratio between the area of the central surface contained in a sphere of a defined size and that of a disk with the same radius (Toro et al., 2008).

### Region-based Processing and Morphometry

In addition to voxel- or point-wise analyses via VBM or SBM, CAT provides an option to conduct regional analyses via region-based morphometry (RBM). For this purpose, the processing steps under voxel-based processing (surface-based processing, respectively) should be applied and followed by automatically calculating regional measurements. This is achieved by working with regions of interest (ROIs), defined using standardized atlases. The required atlases are provided in CAT (see **Supplemental Table 1** and **Supplemental Table 2**), but users can also work with their own atlases.

⍰ Voxel-based ROIs: The volumetric atlases available in CAT have been defined on brain templates in MNI space and may be mapped to the individual brains by using the spatial registration parameters determined during voxel-based processing. Volumetric measures, such as regional gray matter volume, can then be calculated for each ROI in native space.
⍰ Surface-based ROIs: The surface atlases available in CAT are supplied on the *FsAverage* surface and can be mapped to the individual surfaces by using the spherical registration parameters determined during the surface-based processing. Surface-based measures, such as cortical thickness or cortical folding, are then calculated for each ROI in native space.

### Performance of CAT

CAT allows processing streams to be distributed to multiple processing cores, to reduce processing time. For example, CAT12’s analysis of 50 subjects (see **<u>Example Application</u>**) leveraging the inbuilt parallel processing capabilities on four cores, required seven hours processing time when analyzing one image per subject (cross-sectional stream), and 18 hours when processing three images per subject (longitudinal stream) for the entire sample. Application of all available workflows for a single T1-weighted image takes around 35 minutes, as timed on an iMac with Intel Core i7 with 4 GHz and 32 GB RAM using Matlab 2017b, SPM12 r7771, and CAT12.8 r1945.

CAT’s performance has been thoroughly tested by evaluating its accuracy, sensitivity and robustness in comparison to other tools frequently used in the neuroimaging community. For this purpose, we applied CAT12 and analyzed real data (see **<u>Example</u> <u>Application</u>**) as well as simulated data generated from BrainWeb (https://brainweb.bic.mni.mcgill.ca/brainweb). The evaluation procedures are detailed in **<u>Supplemental Note 1</u>** and **<u>Supplemental Note 2</u>**; the outcomes are presented in **Supplemental Figure 1** and **Supplemental Figure 2**. CAT proved to be accurate, sensitive, reliable, and robust outperforming other common neuroimaging tools.

### Five Selected Features of CAT

#### 1. Longitudinal Processing

Aside from offering a standard pipeline for cross-sectional analyses, CAT has specific longitudinal pipelines that ensure a local comparability both across subjects and across time points within subjects. Compared to the cross-sectional pipeline, these longitudinal pipelines render analysis outcomes more accurate when mapping structural changes over time. The user can choose between three different longitudinal pipelines: the first one for analyzing brain plasticity (over days, weeks, months); the second one for analyzing brain development (over months and years); and the third one for brain aging (over months, years, decades). For more details, refer to **<u>Supplemental Note</u> 3.**

#### 2. Quality Control

CAT introduces a retrospective quality control framework for the empirical quantification of essential image parameters, such as noise, intensity inhomogeneities, and image resolution (all of these can be impacted, for example, by motion artifacts). Separate parameter-specific ratings are provided as well as a handy overall rating (Gilmore et al., 2021). Moreover, image outliers can be easily identified, either directly based on the aforementioned indicators of the image quality or by calculating a Z-score determined by the quality of the image processing as well as by the anatomical characteristics of each brain. For more details, refer to **Supplemental Note 4.**

#### 3. Mapping onto the Cortical Surface

CAT allows the user to map voxel-based values (e.g., quantitative, functional, or diffusion parameters) to individual brain surfaces (i.e., pial, central, and/or white matter) for surface-based analyses. The integrated equi-volume model (Bok, 1929) also considers the shift of cytoarchitectonic layers caused by the local folding. Optionally, CAT also allows mapping of voxel values at multiple positions along the surface normal at each node - supporting a layer-specific analysis of ultra-high resolution functional MRI data (Kemper et al., 2017; Waehnert et al., 2013). For more details, refer to **<u>Supplemental Note 5</u>.**

#### 4. Threshold-free Cluster Enhancement (TFCE)

CAT comes with its own TFCE toolbox and provides the option to apply TFCE (Smith & Nichols, 2009) in any statistical *second-level* analysis in SPM, both for voxel-based and for surface-based analyses. It can also be employed to analyze *functional MRI* (fMRI) or *diffusion tensor imaging* (DTI) data. A particularly helpful feature of the TFCE toolbox is that it automatically recognizes exchangeability blocks and potential nuisance parameters (Winkler et al., 2014) from an existing statistical design in SPM. For more details, refer to **<u>Supplemental Note 4</u>.**

#### 5. Visualization

CAT allows a user to generate graphs and images, which creates a solid basis to explore findings as well as to generate ready-to-publish figures according to prevailing standards. More specifically, it includes two distinct sets of tools to visualize results: the first set prepares both voxel- and surface-based data for visualization by providing options for thresholding the default SPM *T*-maps or *F*-maps and for converting statistical parameters (e.g., *T*-maps and *F*-maps into *p*-maps). The second set of tools visualizes the data offering the user ample options to select from different brain templates, views, slices, significance parameters, significance thresholds, color schemes, etc. (see **Figure 5**).

**Figure 5:**
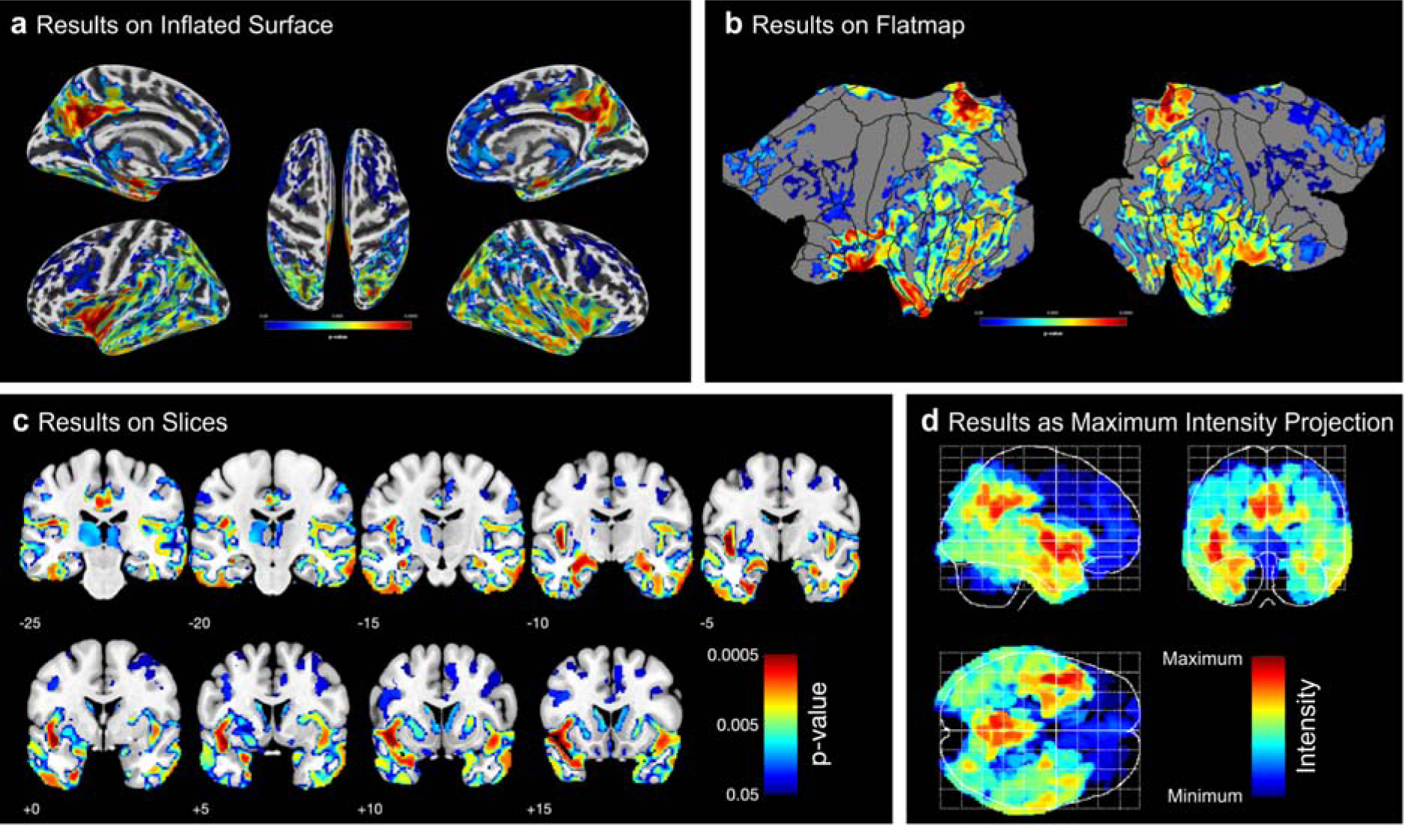
Examples of CAT’s visualization of results. Both surface- and voxel-based data can be presented on surfaces such as (a) the (inflated) FsAverage surface, or (b) the flatmap of the Connectome Workbench. Volumetric maps can also be displayed as (c) slice overlays on the MNI average brain, or (d) as a maximum intensity projection (so-called “glass brains”). All panels show the corrected *p*-values from the longitudinal VBM study in our example (see Example Application).

### Example Application

To demonstrate an application of CAT, we investigated an actual dataset focusing on the effects of Alzheimer’s disease on brain structure. More specifically, we set out to compare 25 patients with Alzheimer’s disease and 25 matched controls. We applied (I) a VBM analysis focusing on voxel-wise gray matter volume, (II) an RBM analysis focussing on regional gray matter volume (i.e., a voxel-based ROI analysis), (III) a surface-based analysis focusing on point-wise cortical thickness, and (IV) an RBM analysis focussing on regional cortical thickness (i.e., a surface-based ROI analysis). Given the wealth of literature on Alzheimer’s disease, we expected atrophy in gray matter volume and cortical thickness in patients compared to controls, particularly in regions around the medial temporal lobe and the default mode network (Bayram et al., 2018; Dickerson, 2010). In addition to distinguishing between the four morphological measures (I-IV), all analyses were conducted using both cross-sectional and longitudinal streams in CAT. Overall, we expected that longitudinal changes would manifest in similar brain regions to cross-sectional group differences, but that cross-sectional effects would be more pronounced than longitudinal effects. The outcomes of this example analysis are presented and discussed in the next section.

## Discussion

### Example Application

As shown in **Figure 6**, all four cross-sectional streams – investigating voxel-based gray matter volume, regional gray matter volume, point-wise thickness, and regional thickness – revealed widespread group differences between AD patients and matched controls. Overall, the effects were comparable between cross-sectional and longitudinal streams, but the significant clusters were more pronounced cross-sectionally (note the different thresholds cross-sectionally and longitudinally).

**Figure 6:**
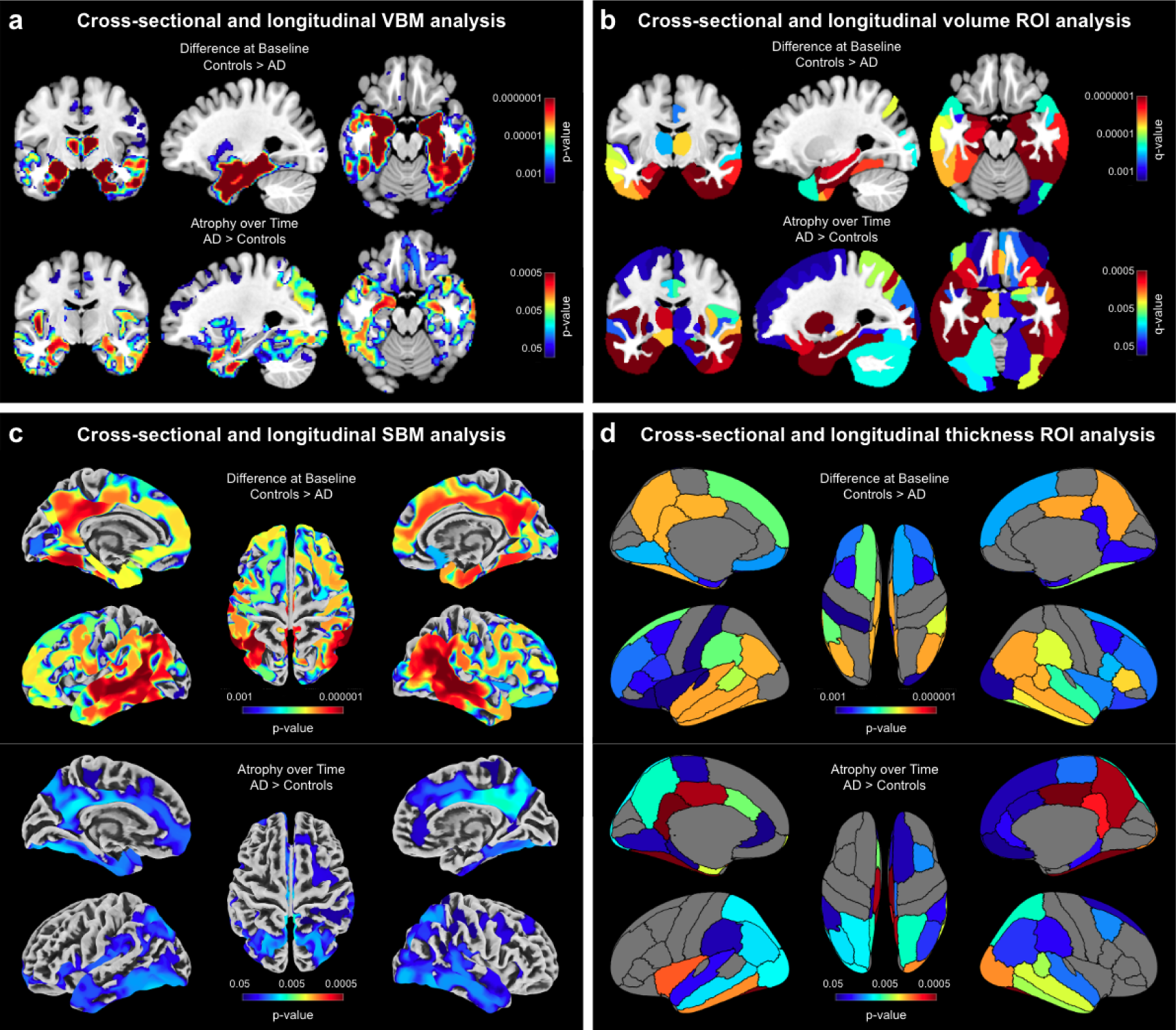
Pronounced atrophy in gray matter and cortical thickness in patients with Alzheimer’s disease compared to healthy control subjects. (a) *Voxel-based Morphometry* (VBM) findings: Results were estimated using *threshold-free cluster enhancement* (TFCE), corrected for multiple comparisons by controlling the *family-wise error* (FWE), and thresholded at p<0.001 for cross-sectional data and p<0.05 for longitudinal data. Significant findings wer projected onto orthogonal sections intersecting at (x=-27mm, y=-10mm, z=-19mm) of the mean brain created from the entire study sample (n=50). (b) Volumetric *Regions of Interest* (ROI) findings: ROIs were defined using the Neuromorphometrics atlas. Results were corrected for multiple comparisons by controlling the *false discovery rate* (FDR) and thresholded at q<0.001 for cross-sectional data and q<0.05 for longitudinal data. Significant findings were projected onto the same orthogonal sections as for the VBM findings. (c) *Surface-based Morphometry* (SBM) findings: Results were estimated using TFCE, FWE-corrected, and thresholded at p<0.001 for cross-sectional data and p<0.05 for longitudinal data. Significant findings wer projected onto the FreeSurfer *FsAverage* surface. (d) Surface *Regions of Interest* (ROI) findings: ROIs were defined using the DK40 atlas. Results were FDR-corrected and thresholded at q<0.001 for cross-sectional data and q<0.05 for longitudinal data. Significant findings were projected onto the *FsAverage* surface.

More specifically, using VBM, significantly smaller voxel-wise gray matter volumes were observed in AD patients compared to controls, particularly in the medial and lateral temporal lobes and within regions of the default mode network (**Figure 6a <u>top</u>**). Similarly, the longitudinal follow-up revealed a significantly stronger gray matter volume loss in patients compared to controls, with effects located in the medial temporal lobe as well as the default mode network (**Figure 6a <u>bottom</u>**). The voxel-based ROI analysis resulted in a significance pattern similar to the VBM study, with particularly pronounced group differences in the temporal lobe that extended into additional brain areas including those comprising the default mode network (**Figure 6b <u>top</u>**). Again, the longitudinal analysis yielded similar but less pronounced findings than the cross-sectional analysis, although longitudinal effects were stronger than in the VBM analysis (**Figure 6b <u>bottom</u>**).

Using SBM, the point-wise cortical thickness analysis yielded a pattern similar to the VBM analysis with significantly thinner cortices in patients, particularly in the medial and lateral temporal lobe and within regions of the default mode network (**Figure 6c <u>top</u>**). Just as in the VBM analysis, significant clusters were widespread and reached far into adjacent regions. Again, the results from the longitudinal stream were less widespread and significant than the results from the cross-sectional stream (**Figure 6c <u>bottom</u>**). Finally, the surface-based ROI analysis largely replicated the local findings from the SBM analysis (**Figure 6d <u>top / bottom</u>**).

Overall, the results of all analysis streams corroborate prior findings in the Alzheimer’s disease literature, particularly the strong disease effects within the medial temporal lobe and regions of the default mode network (Bayram et al., 2018; Dickerson, 2010). Furthermore, the comparable pattern across measures suggests a considerable consistency between available morphometric options, even if gray matter volume and cortical thickness are biologically different and not perfectly related (Hutton et al., 2009; Winkler et al., 2018).

### Evaluation of CAT12

As shown in **Supplemental Figure 1** and **Supplemental Figure 2**, CAT12 proved to be accurate, sensitive, reliable, and robust outperforming other common neuroimaging tools. Similar conclusions have been drawn in independent evaluations testing one or more softwares in comparison with CAT12. For example, Guo et al. (2018) evaluated the repeatability and reproducibility of brain volume measurements using FreeSurfer, FSL-SIENAX and SPM, and highlighted the reliability of CAT12. Similarly, CAT12 emerged as a robust option when demonstrating that the choice of the processing pipeline influences the location of neuroanatomical brain markers (Zhou et al., 2022). Last but not least, Khlif et al. (2019) compared the outcomes of CAT12’s automated segmentation of the hippocampus with those achieved based on manual tracing and demonstrated that both approaches produced comparable hippocampal volume.

In addition, numerous evaluations suggest that CAT12 performs at least as well as other common neuroimaging tools and, as such, offers a valuable alternative. For example, Tavares et al. (2019) conducted a VBM study and concluded that the segmentation pipelines implemented in CAT12 and SPM12 provided results that are highly correlated and that the choice of the pipeline had no impact on the accuracy of any brain volume measure. Along the same lines, but for SBM, Ay et al. (2022) reported that CAT12 and FreeSurfer produced equally valid results for parcel-based cortical thickness calculations. de Fátima Machado Dias et al. (2022) addressed the issue of reproducibility and observed that cortical thickness measures using CAT12 and FreeSurfer were comparable at the individual level. Moreover, Seiger et al. (2018) conducted a study in patients with Alzheimer’s disease and healthy controls, in which CAT12 and FreeSurfer provided consistent cortical thickness estimates and excellent test-retest variability scores. Velázquez et al. (2021) supported these findings when comparing CAT12 and FreeSurfer with three voxel-based methods in a test-retest analysis and clinical application. Finally, Righart et al (2017) compared volume and surface-based cortical thickness measurements in multiple sclerosis and emphasized CAT12’s consistent performance.

These collective findings from multiple studies support the notion that CAT12 is a robust and reliable tool for both VBM and SBM analyses, producing results that are comparable to, and in some cases, superior to, other established neuroimaging softwares.

## Conclusion

CAT is suitable for desktop and laptop computers as well as high-performance clusters. It is fully integrated into the SPM environment within Matlab, but also allows process execution directly from the command shell, without having to start SPM. CAT is also available as a standalone version, avoiding the need for a Matlab license. In terms of performance, CAT allows for ultra-fast processing and analysis and also is more sensitive in detecting significant effects compared to other common tools used by the neuroimaging community. Moreover it better handles varying levels of noise and signal inhomogeneities. Furthermore, CAT is easy to integrate with non-SPM software packages and also supports the Brain Imaging Data Structure (BIDS) standards (Krzysztof et al., 2015). Therefore, CAT is ideally suited not only to process small datasets (as demonstrated in the example application), but also big datasets, such as samples of the UK Biobank (https://www.ukbiobank.ac.uk) or ENIGMA (https://enigma.ini.usc.edu). Finally, while CAT is currently targeted at structural imaging data, some features (e.g., high-dimensional spatial registration or mapping onto the cortical surface) may also be used for the analysis of functional, diffusion, or quantitative MRI or EEG/MEG data.

## Methods

### Application Example

#### Data Source

Data for the application example were obtained from the Alzheimer’s Disease Neuroimaging Initiative (ADNI) database (<u>adni.loni.usc.edu</u>). The ADNI was launched in 2003 as a public-private partnership, led by Principal Investigator Michael W. Weiner, MD. The primary goal of ADNI has been to test whether serial magnetic resonance imaging (MRI), positron emission tomography (PET), other biological markers, and clinical and neuropsychological assessment can be combined to measure the progression of mild cognitive impairment (MCI) and early Alzheimer’s disease (AD). For up-to-date information, see www.adni-info.org.

#### Sample Characteristics

For the purpose of the current study, we compiled a sample of fifty subjects with 3D T1-weighted structural brain images from the ADNI database. Specifically, we randomly selected the first 25 subjects (16 males / 9 females) classified as AD patients (mean age 75.74±8.14 years; mean *minimal mental status examination* (MMSE) score: 23.44±2.04) and matched them for sex and age with 25 healthy controls (mean age 76.29±3.90 years; mean MMSE: 28.96±1.24). Informed consent was obtained from all research participants. All subjects had brain scans at baseline (first scan at enrolment) and at two follow-up visits, at one year and at two years after the first scan. All brain images were acquired on 1.5 Tesla scanners (Siemens, General Electric, Philips) using a 3D T1-weighted sequence with an in-plane resolution between 0.94 and 1.25 mm and a slice thickness of 1.2 mm.

#### Data Processing

All T1-weighted data were processed using CAT12 following the cross-sectional (or longitudinal, respectively) processing stream for VBM, SBM (cortical thickness), and ROI analyses (see **Figure 2**) according to the descriptions provided under **Computational Morphometry**. For each subject, only their first time point was included in the cross-sectional stream, whereas all three time points were included in the longitudinal stream. The processing streams for the VBM analysis resulted in modulated and registered gray matter segments, which were smoothed using a 6 mm Gaussian kernel. The image processing streams for the SBM analysis resulted in the registered point-wise cortical thickness measures, which were smoothed using a 12 mm Gaussian Kernel. The voxel-based ROI analysis used the Neuromorphometrics atlas (http://Neuromorphometrics.com/) to calculate the regional gray matter volumes; the surface-based ROI analysis employed the DK40 atlas (Desikan et al., 2006) to calculate regional cortical thickness.

#### Statistical Analysis

For each variable of interest – voxel-wise gray matter volume, regional gray matter volume, point-wise cortical thickness, and regional cortical thickness – the dependent measures (e.g., the registered, modulated, and smoothed gray matter segments for voxel-wise gray matter) were entered into the statistical model. For the cross-sectional stream, *group* (Alzheimer’s disease patients vs. controls) was defined as the independent variable. For the longitudinal stream, the interaction between *group* and *time* was defined as the independent variable, whereas *subject* was defined as a variable of no interest. For the VBM and the voxel-based ROI analyses, data were corrected for TIV using ‘global scaling’ (because TIV correlated with *group*, the effect of interest). Since cortical thickness does not scale with brain size (Barnes et al., 2010), no corrections for TIV were applied for the SBM and the surface-based ROI analyses. For the cross-sectional analysis we additionally included age as a nuisance parameter.

For the VBM and SBM analyses, results were corrected for multiple comparisons by applying TFCE (Smith & Nichols, 2009) and controlling the family-wise error at p≤0.001 (cross-sectional) and *p*≤0.05 (longitudinal). For the voxel-based and surface-based ROI analyses, results were corrected for multiple comparisons by controlling the false discovery rate (Benjamini & Hochberg, 1995) at *q*≤0.001 (cross-sectional) and *q*≤0.05 (longitudinal). All statistical tests were one-tailed given our a priori hypothesis that AD patients present with less gray matter at baseline and a larger loss of gray matter over time.

The outcomes of the VBM and voxel-based ROI analyses were overlaid onto orthogonal sections of the mean brain created from the entire study sample (n=50); the outcomes of the SBM and surface-based ROI analyses were projected onto the *FsAverage* surface.

## Acknowledgments

This article received support through the Humboldt Foundation (Germany), the Royal Society of New Zealand (20-UOA-045), a Hood Fellowship from the University of Auckland (New Zealand), the Federal Ministry of Science and Education (BMBF) under the frame of ERA PerMed (Pattern-Cog ERAPERMED2021-127), the Marie Skłodowska-Curie Innovative Training Network (SmartAge 859890 H2020-MSCA-ITN2019), and the Carl Zeiss Foundation (IMPULS P2019-01-006). It was also supported by the Eunice Kennedy Shriver National Institute of Child Health & Human Development of the National Institutes of Health (R01HD081720) and by the Royal Society of New Zealand (Marsden 20-UOA-045). Data collection and sharing for this project was funded by the Alzheimer’s Disease Neuroimaging Initiative (ADNI) (National Institutes of Health Grant U01 AG024904) and DOD ADNI (Department of Defense award number W81XWH-12-2-0012). ADNI is funded by the National Institute on Aging, the National Institute of Biomedical Imaging and Bioengineering, and through generous contributions from the following: AbbVie, Alzheimer’s Association; Alzheimer’s Drug Discovery Foundation; Araclon Biotech; BioClinica, Inc.; Biogen; Bristol-Myers Squibb Company; CereSpir, Inc.; Cogstate; Eisai Inc.; Elan Pharmaceuticals, Inc.; Eli Lilly and Company; EuroImmun; F. Hoffmann-La Roche Ltd and its affiliated company Genentech, Inc.; Fujirebio; GE Healthcare; IXICO Ltd.; Janssen Alzheimer Immunotherapy Research & Development, LLC.; Johnson & Johnson Pharmaceutical Research & Development LLC.; Lumosity; Lundbeck; Merck & Co., Inc.; Meso Scale Diagnostics, LLC.; NeuroRx Research; Neurotrack Technologies; Novartis Pharmaceuticals Corporation; Pfizer Inc.; Piramal Imaging; Servier; Takeda Pharmaceutical Company; and Transition Therapeutics. The Canadian Institutes of Health Research is providing funds to support ADNI clinical sites in Canada. Private sector contributions are facilitated by the Foundation for the National Institutes of Health (www.fnih.org). The grantee organization is the Northern California Institute for Research and Education, and the study is coordinated by the Alzheimer’s Therapeutic Research Institute at the University of Southern California. ADNI data are disseminated by the Laboratory of Neuro Imaging (LONI) at the University of Southern California (USC).

## Data Availability

MRI data are available after obtaining approval to access ADNI data at https://adni.loni.usc.edu. The BrainWeb data is available at https://brainweb.bic.mni.mcgill.ca/.

## Code Availability

The Computational Anatomy Toolbox is available under the GPL license at https://github.com/ChristianGaser/cat12. All code used to perform the statistical tests are available at https://github.com/ChristianGaser/tfce, under the GPL License. Software documentation is available at https://neuro-jena.github.io/cat12-help.

## Supplemental Material

### Supplemental Notes

#### Supplemental Note 1: Comparison with other tools

We evaluated the performance of CAT12 by comparing it to other tools commonly used in the neuroimaging community. More specifically, we assessed the accuracy and sensitivity using CAT12, SPM12, FSL-FAST6, Freesurfer6 and CIVET 2.1 in detecting subtle alterations in brain structure that are critical for early diagnosis and monitoring of Alzheimer’s disease. Note, the primary aim of our comparison is to provide insights into the tool’s performances; revealing aberrations associated with Alzheimer’s disease is only a secondary aim of this paper. To conduct the comparisons we used the same baseline data of our example application (25 patients with Alzheimer’s disease and 25 matched controls), as described in the main article. The analyses focussed on (1) voxel-wise gray matter volume and (2) point-wise cortical thickness. Analyses pertaining to (1) were conducted using voxel-based morphometry (VBM) while processing the data with (1a) SPM12 (https://www.fil.ion.ucl.ac.uk/spm) as well as with (1b) FSL-FAST6 (https://www.fmrib.ox.ac.uk/fsl). Analyses pertaining to (2) were conducted using surface-based morphometry (SBM) while processing the data with (2a) Freesurfer7.2 (https://surfer.nmr.mgh.harvard.edu) as well as with (2b) Civet2.1 (https://mcin.ca/technology/civet).

#### Data Processing for VBM data

1a – SPM12: We applied the Unified Segmentation (Ashburner & Friston, 2005)(Ashburner and Friston, 2005) in SPM12 with default settings to extract rigidly registered gray and white matter segments. These individual segments provided the basis to create a mean segment using the Shooting toolbox (Ashburner & Friston, 2011) in SPM12. This mean segment functions as an initial template and is warped to each of the individual segments, which is followed by calculating the resulting deformations, applying the inverses of the deformations to the individual images, and re-calculating the template (aka the mean segment). This process is repeated several times. The results are spatially registered segments which will be adjusted for volume changes introduced by the registration (modulation) and convoluted with a Gaussian kernel of FWHM 6mm (smoothing).

1b – FSL-FAST6: We applied the FSLVBM script from FSL6 to process the data (https://fsl.fmrib.ox.ac.uk/fsl/fslwiki/FSLVBM/UserGuide). The default there is using BET to skull-strip the data. However, the achieved output was of poor quality, which is why we used the aforementioned SPM12 segments (in native space) to skull-strip the data. The skull-stripped data were then processed using the FSLVBM script and smoothed with a 6mm Gaussian kernel, as described above.

#### Data Processing for SBM data

2a – Freesurfer7.2 The data were processed using the recon-all script for Freesurfer7.2 (https://surfer.nmr.mgh.harvard.edu) with default settings. For a better comparison between tools, the resulting cortical thickness measures were resampled and smoothed (FWHM 12 mm) using CAT12.

2b – CIVET2.1: The data were uploaded to CBRAIN (https://portal.cbrain.mcgill.ca) and processed with the CIVET2.1 pipeline (https://mcin.ca/technology/civet) using default settings. Again, the cortical thickness measures were resampled and smoothed (FWHM 12 mm) using CAT to allow for a better comparison between tools.

#### Statistical Analysis

For details on the statistical model (e.g., dependent variables, independent variables, and variables of no interest), refer to the Methods section in the main document. All results were corrected for multiple comparisons by applying TFCE (Smith & Nichols, 2009) and controlling the family-wise error at *p*<0.001. All statistical tests were one-tailed given our a priori hypothesis that AD patients present with less gray matter at baseline and a larger loss of gray matter over time. In addition, we calculated the effect sizes to allow for a direct comparison across tools with respect to their sensitivity in detecting significant differences between AD patients and controls.

#### Supplemental Note 2: Evaluation with simulated data

To comprehensively evaluate the performance of CAT12 in comparison with other neuroimaging tools (SPM12 and FSL-FAST6), we conducted evaluations using simulated data generated from BrainWeb (https://brainweb.bic.mni.mcgill.ca/brainweb). More specifically, we compared the output of CAT12, SPM12, and FSL-FAST6 to ground truth data represented by a brain phantom. As the phantom contains known variations in noise levels and signal inhomogeneities, it aids in objectively assessing the accuracy and robustness of CAT12 and the other tools in dealing with different sources of variation. To measure the agreement between the ground truth and the results of CAT12, SPM12, and FSL-FAST6, we calculated the kappa coefficient.

#### Supplemental Note 3: Longitudinal Processing

The majority of morphometric studies are based on cross-sectional data in which one image is acquired for each subject. Nevertheless, the mapping of structural changes over time requires specific longitudinal designs that consider additional time points (and thus images) for each subject. In theory, all images could be processed using the standard cross-sectional processing workflow. In practice, however, longitudinal data strongly benefit from workflows specifically tailored towards longitudinal analyses, where MR-based noise and inhomogeneities are further reduced and where spatial correspondences are ensured, the latter not only across subjects but also across time points within subjects (Ashburner & Ridgway, 2012; Reuter et al., 2010; Reuter & Fischl, 2011). As a consequence, analyses become more sensitive, as shown in **Supplemental Figure 3**.

CAT12 offers three optimized processing pipelines for longitudinal studies: One for neuroplasticity, one for aging, and one for neurodevelopmental studies. Studies in the framework of neuroplasticity are confined to short time-frames of weeks to months, while studies in the framework of aging and neurodevelopment cover longer time frames of years and, sometimes, even decades. For such extended study durations, it is particularly important to model systematic changes of the brain over time to maintain a voxel- or point-wise comparability across time points. Studies in the framework of neurodevelopment require additional considerations of increasing brain and head sizes. A detailed description of all three longitudinal processing workflows is provided in **Supplemental Figure 4**.

#### Supplemental Note 4. Quality Control

Processing of MRI data strongly depends on the quality of the input data. Multi-center studies and data sharing projects, in particular, need to take into account varying image properties due to different scanners, sequences and protocols. However, even scans acquired on a single scanner and using the same scanning protocol may vary due to motion or other miscellaneous artifacts. CAT12 provides options to perform quality checks, both on the subject level and on the group level. More specifically, on the subject level, CAT12 introduces a novel retrospective quality control framework for the quantification of quality differences between different scans obtained on a single scanner or across different scanners. The quality control allows for the evaluation of essential image parameters (i.e., noise, intensity inhomogeneities, and image resolution) and is automatically performed for each brain when running CAT12’s image processing workflow (see **Supplemental Figure 5**). On the group-level, CAT12 provides options to check and visualize the homogeneity of the entire study sample, thus allowing the user to identify any outliers (see **Supplemental Figure 6**).

#### Supplemental Note 5. Mapping onto the Cortical Surface

Surface-based analyses offer some advantages over voxel-based approaches, such as better inter-subject registration and surface-based smoothing, which may result in a larger statistical power and improved accuracy (Dahnke & Gaser, 2018; Tucholka et al., 2012). CAT12 provides a range of options to map voxel-based values (e.g., functional, quantitative or diffusion parameters) to individual brain surfaces for a subsequent surface-based analysis. For this purpose, voxel-based values are extracted at multiple positions along the surface normal at each node of the surface (see **Supplemental Figure 7**). The exact positions along the surface normal are determined by an equi-volume model (Bok, 1929), which reflects the normal shift of cytoarchitectonic layers caused by the local folding. In addition to default settings, users can specify both the number and location of those positions along the surface normal. The extracted values along the surface normal are then summarized as one value per node. The default here is to summarize values by using the absolute maximum value. However, other options than using the absolute maximum exist, such as using the minimum, mean, or weighted mean value. Alternatively, users may choose to map voxel values at a specified distance (in mm) from the surface or even at multiple position along the surface normal. The latter is useful, for example, when conducting a layer-specific analysis of ultra-high resolution functional MRI data (Kemper et al., 2017; Waehnert et al., 2013).

#### Supplemental Note 6. Threshold-free Cluster Enhancement (TFCE)

SPM’s standard correction for multiple comparisons is based either on the magnitude of the T or F statistic (correction on voxel-level) or on the extent of clusters in a thresholded statistical map (correction on cluster level). The principle of TFCE – as implemented in CAT12’s TFCE toolbox – is to combine both approaches, which has several theoretical and practical advantages, as detailed elsewhere (Smith & Nichols, 2009). Briefly, it retains the sensitivity of cluster-based inferences, while avoiding their main downsides, such as arbitrary cluster-forming thresholds or susceptibility to non-stationarity that may compromise the statistical validity (Eklund et al., 2016; Hayasaka et al., 2004; Salimi-Khorshidi et al., 2010). As a special feature in CAT, the TFCE toolbox automatically recognizes exchangeability blocks and potential nuisance parameters (Winkler et al., 2014), which would otherwise need to be specified by the user.

---

**Supplemental Figure 1:**
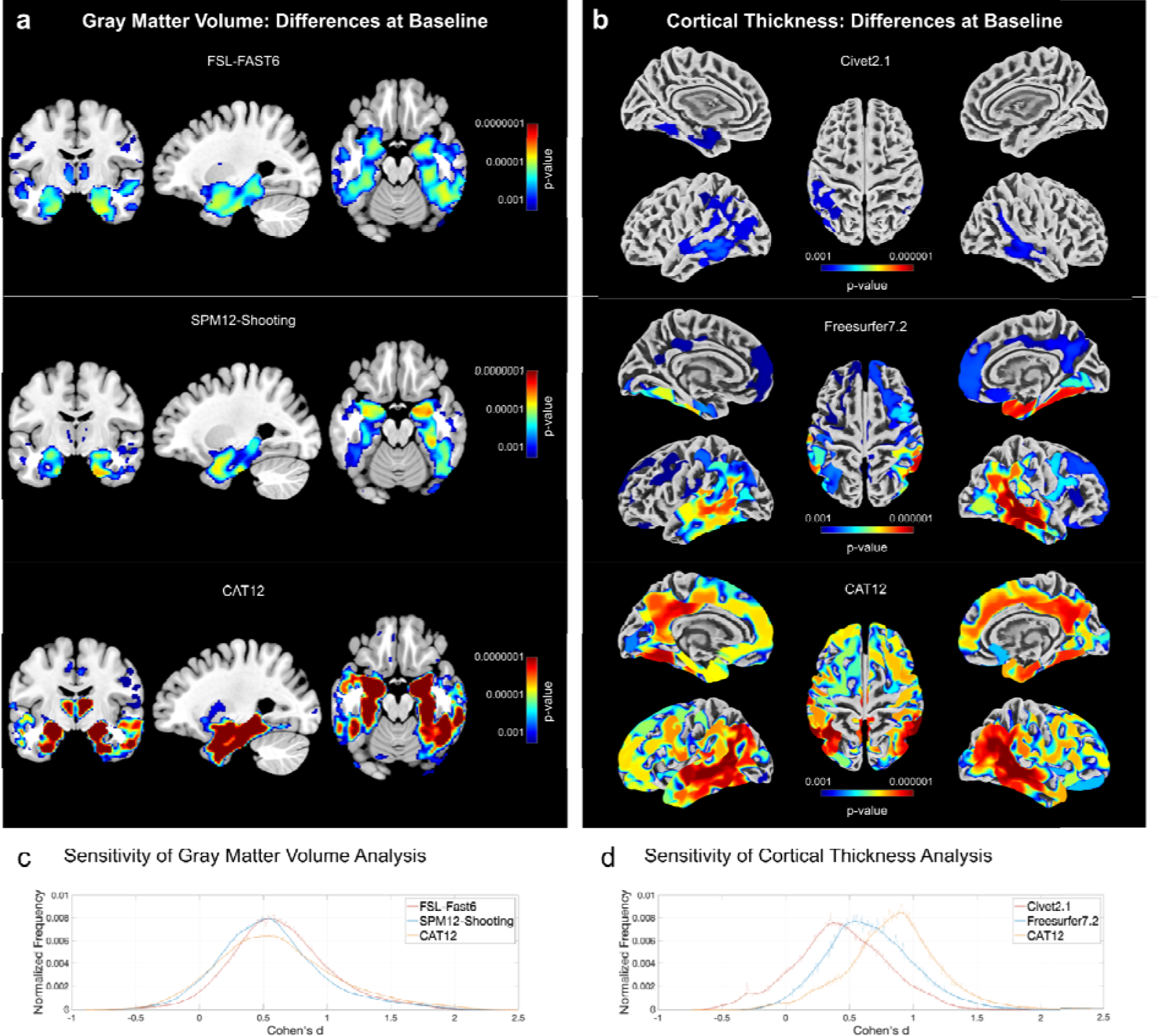
Comparisons between CAT12 and other common tools. *Panel a:* VBM analyses of voxel-wise gray matter volume using FSL-FAST6 (*top*), SPM12-Shooting (*middle*), and CAT12 (*bottom*). *Panel b:* SBM analyses of point-wise cortical thickness using CIVET2.1 (*top*), Freesurfer7.2 (*middle*), and CAT1 (*bottom*). Panels c and d: Sensitivity of VBM and SBM analyses. The effect sizes (Cohen’s *d*) are shown on th *x*-axis; their frequency is shown on the *y*-axis (occurrence is normalized to one to facilitate comparisons between histograms). For both VBM and SBM, CAT12 demonstrates a larger sensitivity in detecting structural differences. This is reflected in the more extended significance clusters and lower *p*-values (*panels a and b*) as well as larger effect sizes (*panels c and d*).

**Supplemental Figure 2:**
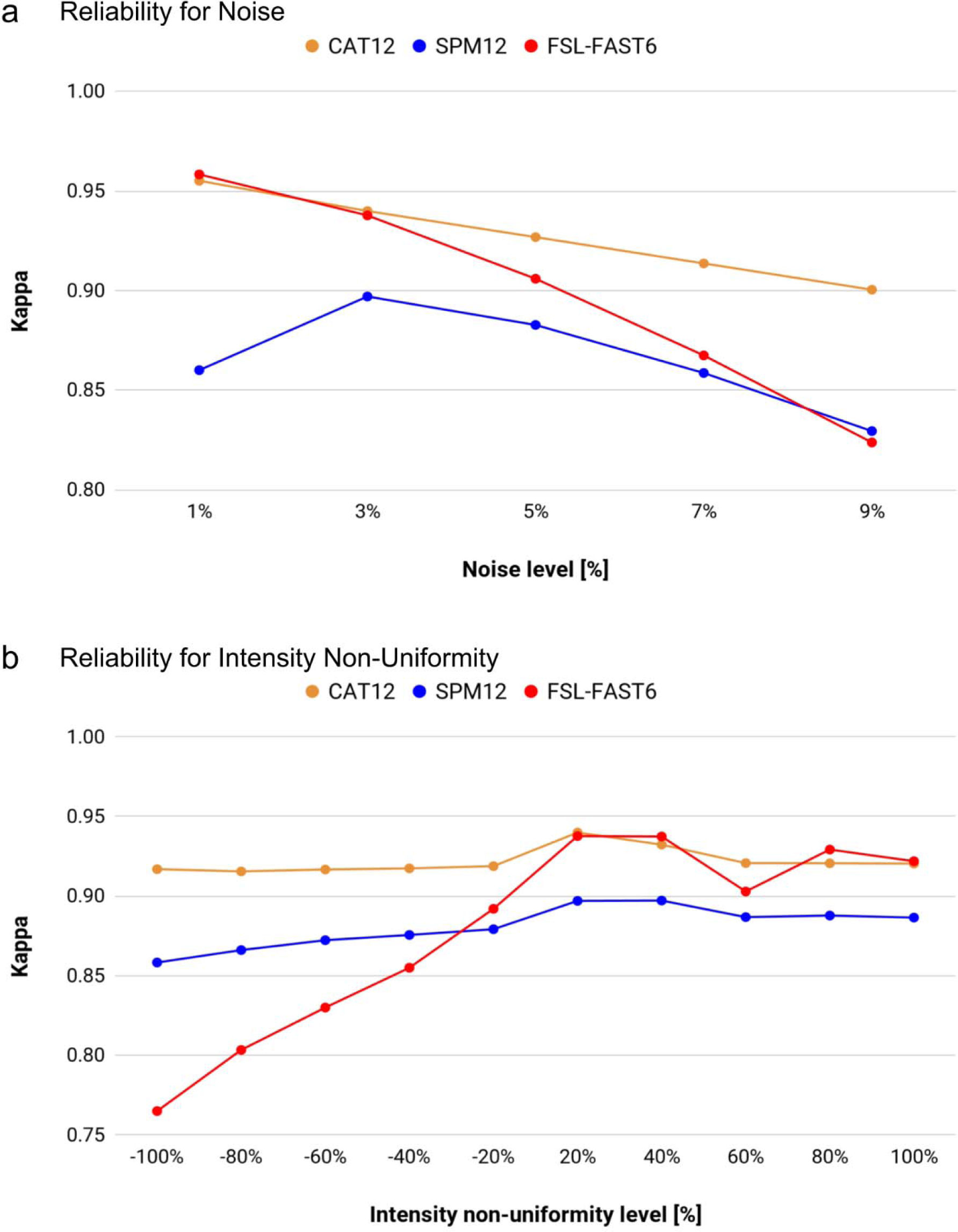
Evaluation of CAT12 and other common tools using Brainweb data. Higher *kapp* values correspond to a better overlap, larger reliability, and increased robustness. *Panel a:* Overlap between ground truth and segmentation outputs for different noise levels. CAT12 is similar to FSL-FAST6 at lower nois levels but clearly outperforms both SPM12 and FSL-FAST6 at higher noise levels. The latter is due to the implemented denoising step (see also Figure 3a for the effect of denosing). *Panel b:* Overlap between ground truth and segmentation outputs for different signal inhomogeneities. CAT12 is extremely robust across th entire range of intensity non-uniformity; it outperforms both SPM12 and FSL-FAST6.

**Supplemental Figure 3:**
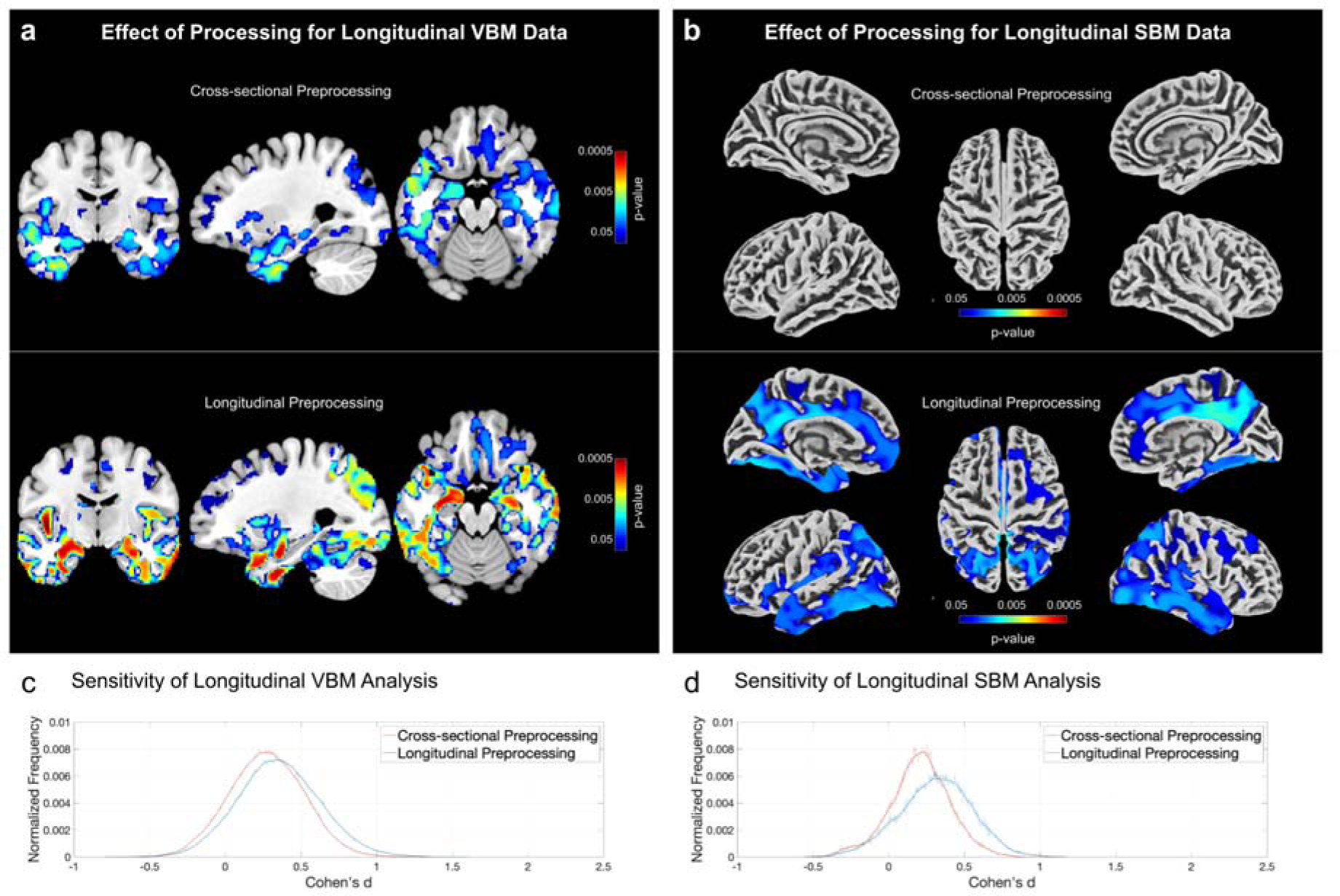
Comparison between CAT12’s cross-sectional and longitudinal pipelines usin longitudinal data. *Voxel-based morphometry* (VBM) results are shown on the left and *surface-based morphometry* (SBM) results on the right. For both VBM and SBM the longitudinal preprocessing leads to an increased sensitivity compared to cross-sectional processing, which is evident as larger clusters and lower p-values (panels a and b) as well as larger effect sizes (panels c and d). The effect sizes are captured as Cohen’s on the x-axis with the frequency of its occurrence normalized to a total sum of one (to ease comparisons between histograms) on the y-axis.

**Supplemental Figure 4:**
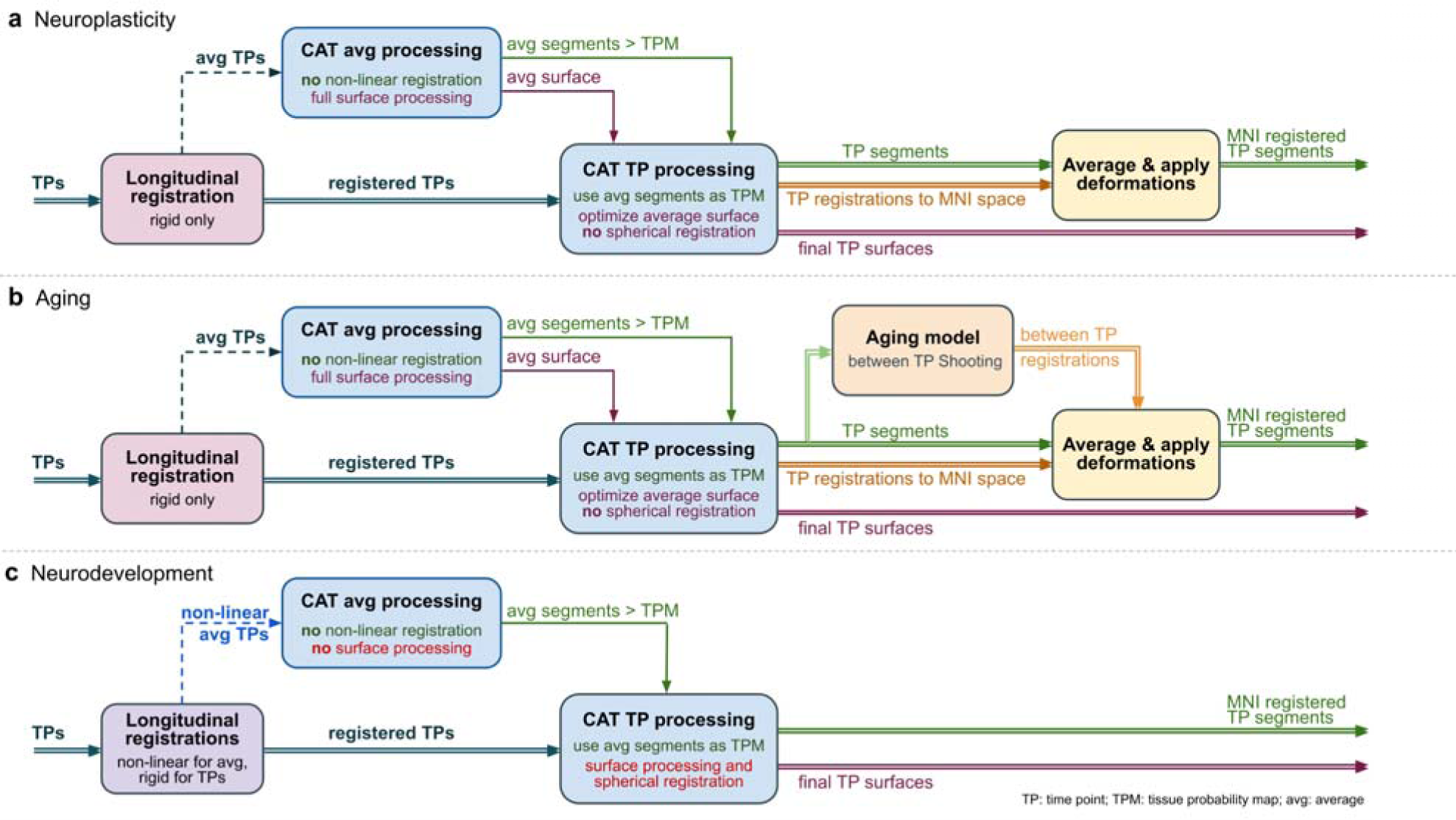
CAT12’s longitudinal processing workflows to examine (a) neuroplasticity, (b) aging, and (c) neurodevelopment. The first step in all three workflows is the creation of a high-quality average imag over all time points. For this purpose, CAT12 realigns the images from all time points for each participant usin inverse-consistent (or symmetric) rigid-body registrations and intra-subject bias field correction. While this is sufficient to create the required average image for the neuroplasticity and aging workflows, the neurodevelopmental workflow requires non-linear registrations in addition. In either case, the resultin average image is segmented using CAT12’s regular processing workflow to create a subject-specific *tissu probability map* (TPM). This TPM is used to enhance the time point-specific processing to create the final segmentations. The final tissue segments are then registered to MNI space to obtain a voxel-comparability across time points and subjects, which differs between all three workflows. In the neuroplasticity workflow, a average of the time point-specific registrations is created to transform the tissue segments of all time points to MNI space. The aging workflow does the same in principle but adds additional (very smooth) deformations between the individual images across time points to account for inevitable age-related changes over time (e.g., enlargements of the ventricles). In contrast, the neurodevelopmental workflow needs to account for major changes, such as overall head and brain growth, which requires independent non-linear registrations to MNI space of all images across time points (which are obtained using the default cross-sectional registration model).

**Supplemental Figure 5:**
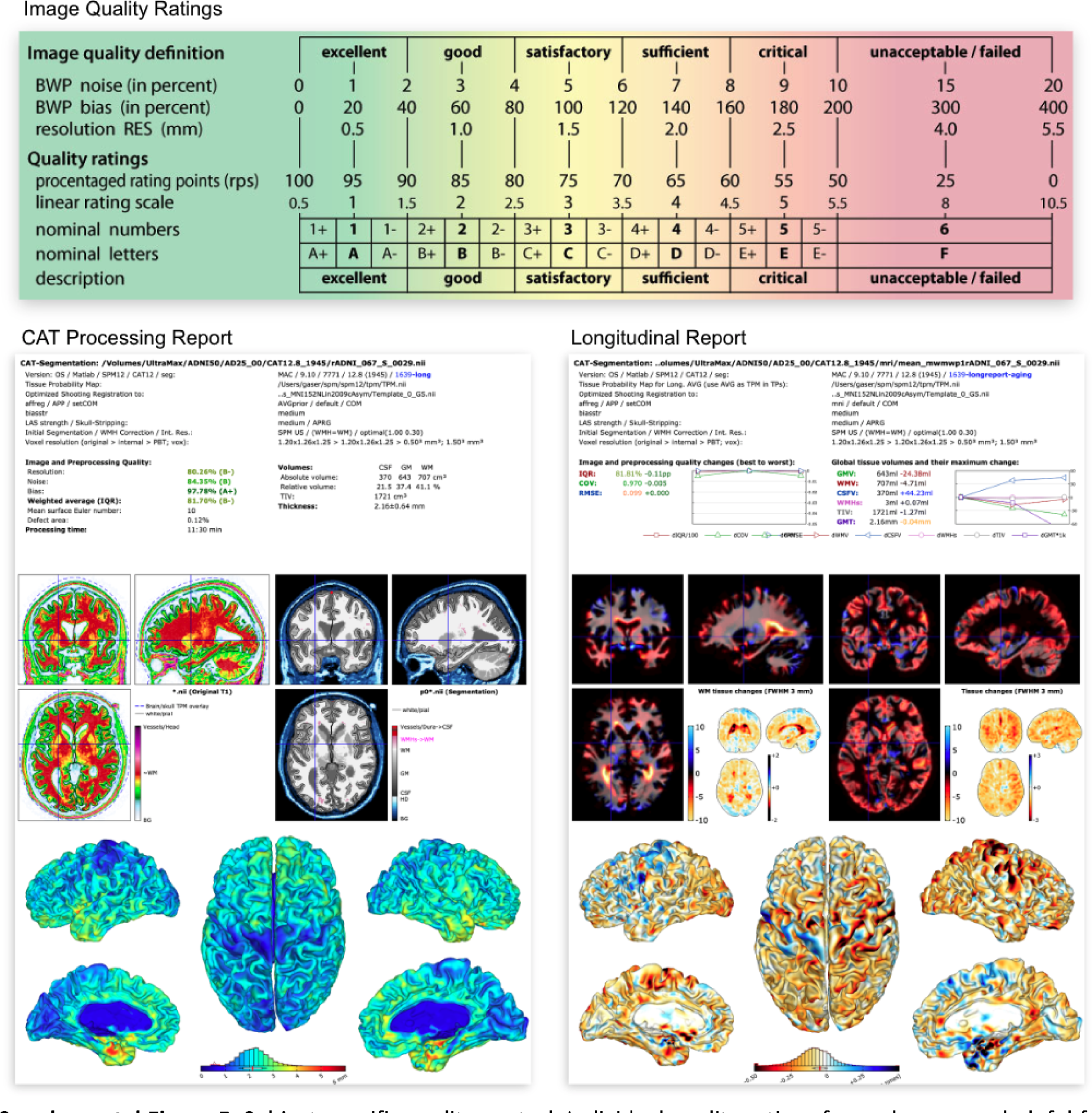
Subject-specific quality control. Individual quality ratings for each scan are helpful for determining potential problems and issues for the use of single scans. The ‘Image Quality Ratings’ (top) employ measures of noise, bias, and image resolution to generate a summary grade for each image (Gilmor et al., 2021). A ‘CAT Processing Report’ (*left*) is automatically saved for each image after the processing workflow is completed; it provides information on image quality measures and the overall grade, in additio to visualizations which allow for an easy assessment of the quality of the skull stripping, tissue segmentation, and surface mapping. Moreover, a ‘Longitudinal Report’ (*right*) is automatically saved when any of the longitudinal pipelines have been used (see **Supplemental Note** 3). This longitudinal report – considering all images of one brain across all time points – provides the same information as the standard cross-sectional report but focuses on the assessment of differences between the individual time points.

**Supplemental Figure 6:**
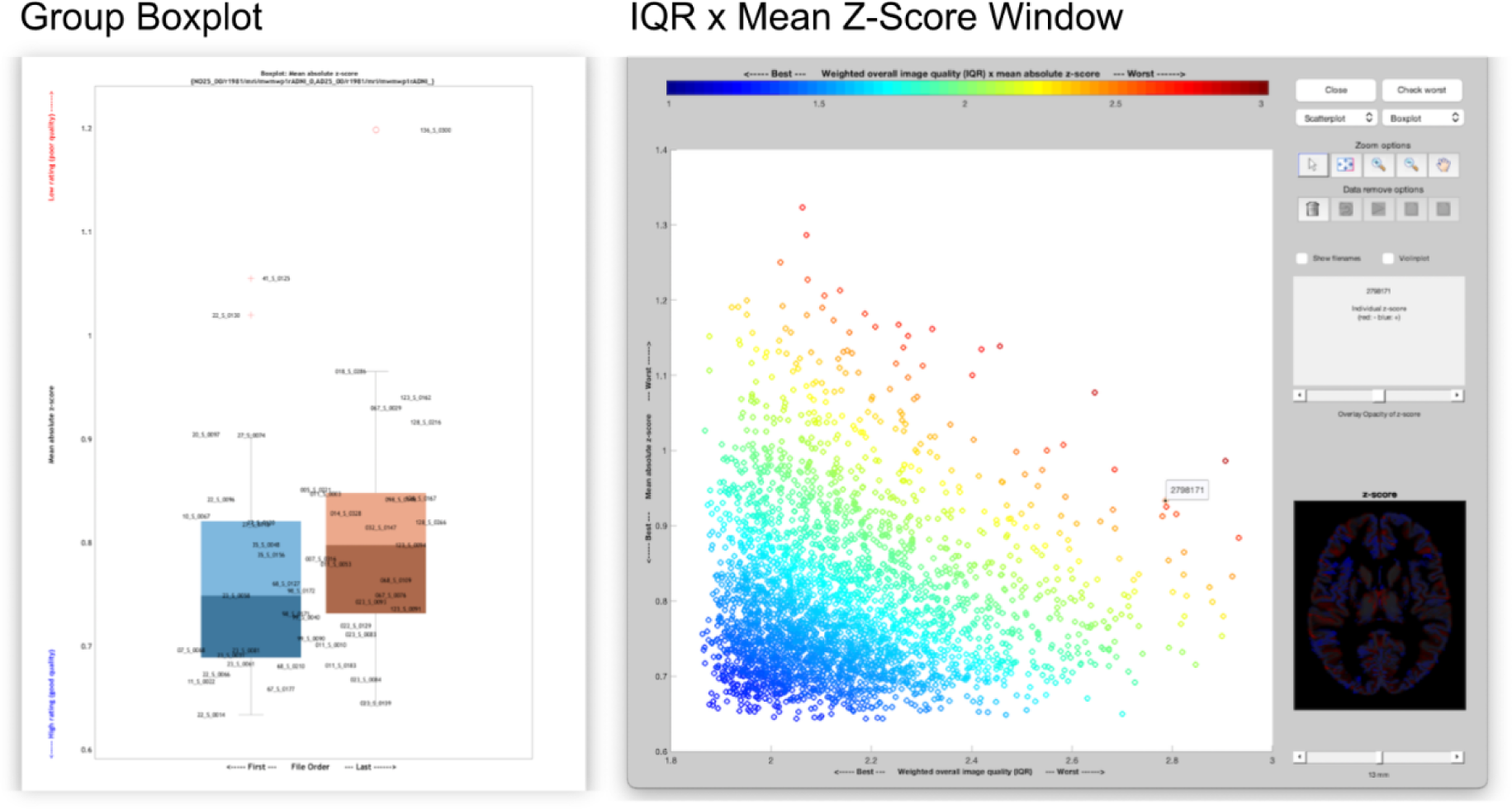
Group-specific quality control. In addition to the subject-specific quality control, larger studies in particular might benefit from scrutinizing those images that are either low in their individual quality ratings and/or different from the other images, suggesting anatomic anomalies, imperfect processing, or other issues that might hamper the subsequent statistical analysis. The ‘Group Boxplot’ (left) allows one to compare any image based on their similarity to the mean and reflects the homogeneity of the sample, by calculatin the average Z-score of all spatially registered images (or surface parameter files). Lower average Z-scor values indicate that the data points are more similar to the mean. Outliers (i.e., images with high Z-scor values) indicate either a potential problem (with the image per se or with the outcomes of the imag processing), or simply a variation in the neuroanatomy (e.g., enlarged ventricles). Such outliers should b checked carefully. An additional ‘IQR x Mean Z-Score Window’ (right) compares the average Z-scores with the weighted image quality rating (IQR) for each subject and allows a combined view of sample homogeneity an overall image quality.

**Supplemental Figure 7:**
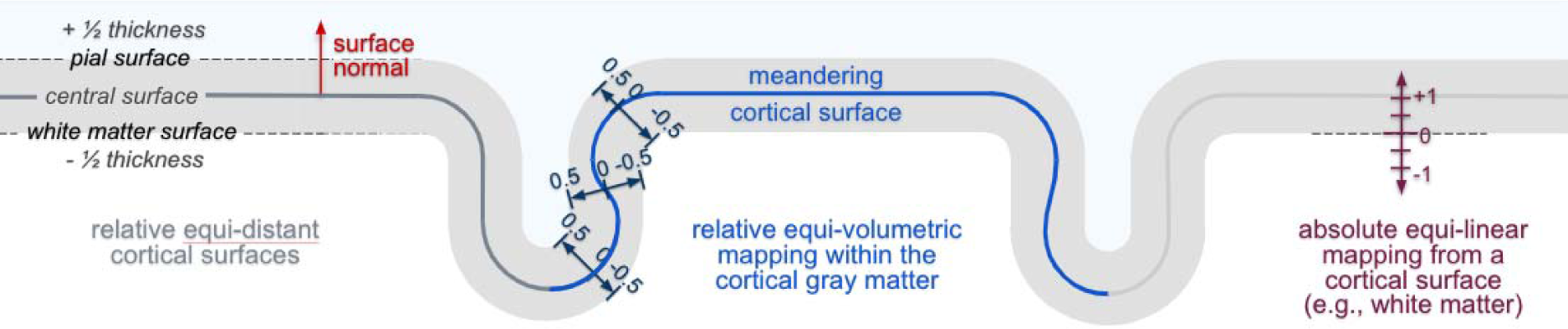
Volume mapping. CAT12 offers multiple ways to map voxel values onto the surface. The default mapping extracts voxel values at multiple positions along a surface normal between the whit matter surface and the pial surface. The exact location of these positions along the normal is determined by an equi-volumetric model (Bok, 1929), which reflects the shift of cortical layers caused by local folding. However, voxel values can also be extracted at a specific user-defined displacement (in mm) from any given surface location.

## Supplemental Tables

**Supplemental Table 1:**
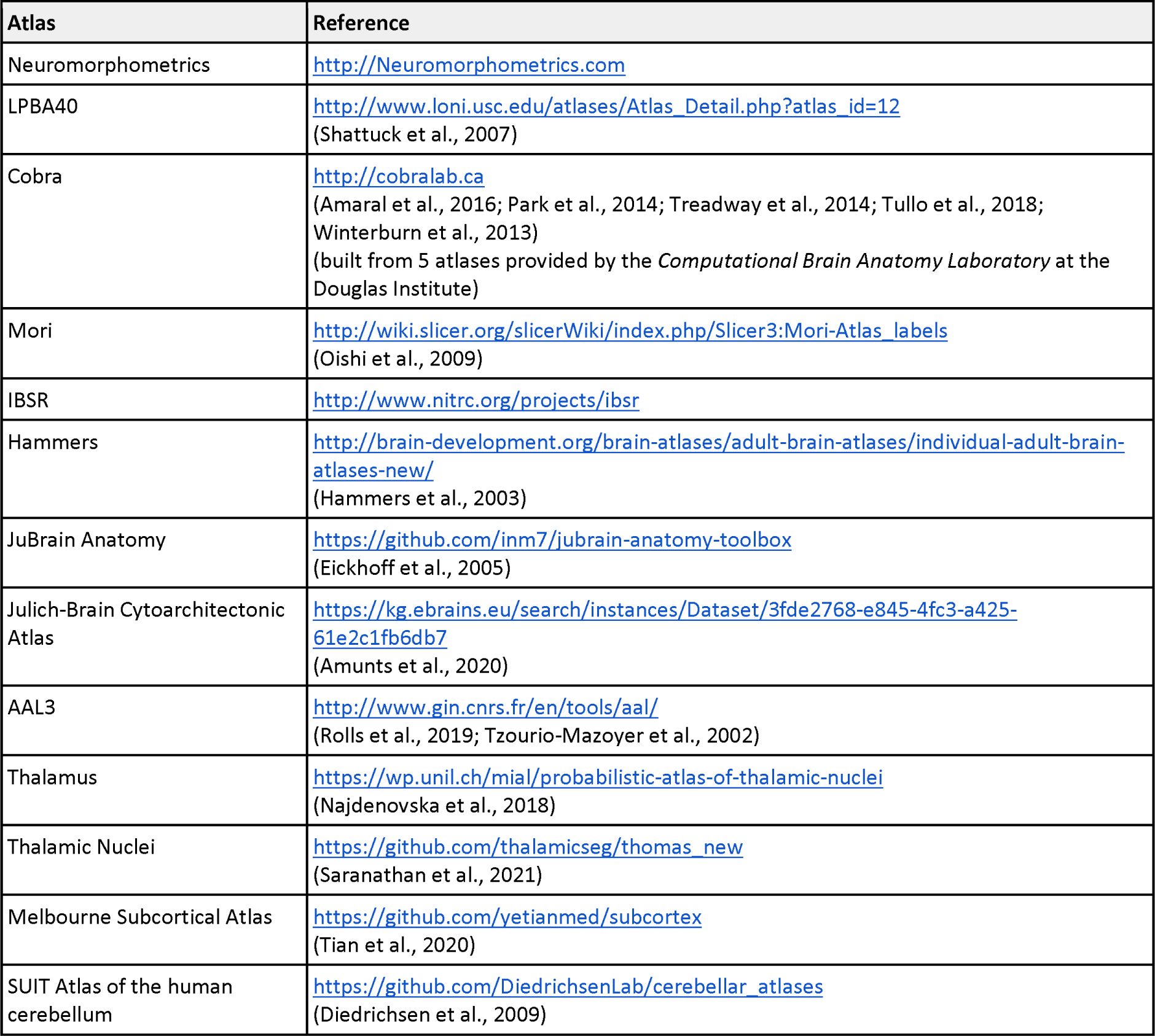
Voxel-based ROI atlases available in CAT12 (as of October 2023)

**Supplemental Table 2:**
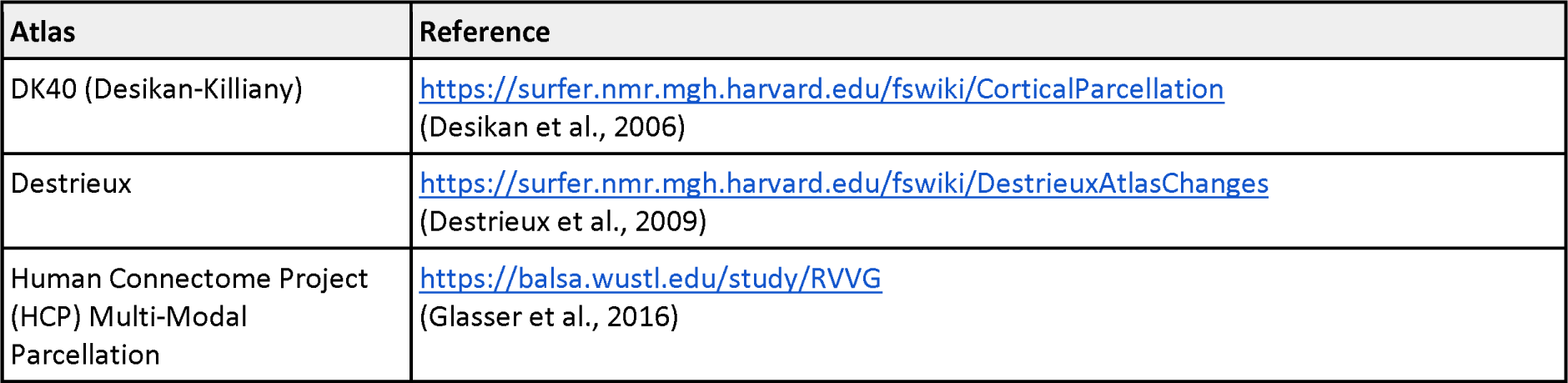
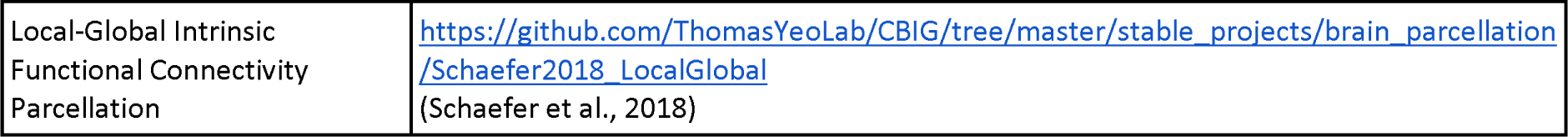
Surface-based ROI atlases available in CAT (as of October 2023)

